# CRISPR-Cas9 mutated pregnane x receptor (*pxr*) retains pregnenolone-induced expression of cytochrome p450 family 3, subfamily A, polypeptide 65 (*cyp3a65*) in zebrafish (*Danio rerio*) larvae

**DOI:** 10.1101/652743

**Authors:** Matthew C. Salanga, Nadja R. Brun, Rene Francolini, John J. Stegeman, Jared V. Goldstone

## Abstract

Pregnane x receptor *(PXR)* is a nuclear receptor that regulates transcriptional responses to drug or xenobiotic exposure in many vertebrate species. One key response is the induction of cytochrome P450 3A *(CYP3A)* transcription. PXR is a promiscuous receptor activated by a wide range of ligands that can differ across species, making functional studies on its role in the chemical defensome, most relevant when approached in a species-specific manner. Genetic knockout studies in mammals have shown a requirement for PXR in ligand-dependent activation of *CYP3A* expression or reporter gene activity. Transient knockdown in zebrafish revealed a similar requirement, however it is not known what the effect of a genetic knockout would be in the zebrafish model. Here, we report on generation of two zebrafish lines each carrying a genetic deletion in the *pxr* coding region, predicted to result in loss of a functional gene product. To our surprise zebrafish larvae, homozygous for either of the *pxr* mutant alleles, retain their ability to induce *cyp3a65* mRNA expression following exposure to the established zebrafish Pxr ligand, pregnenolone (PN). Thus, zebrafish carrying *pxr* alleles with sizable deletions in either the DNA binding or the ligand binding domains do not yield a loss-of-function phenotype, suggesting that a compensatory mechanism is responsible for *cyp3a65* induction. Two alternative possibilities are that Pxr is not required for the effect or that truncated yet functional mutant Pxr is sufficient for the effect.

## Introduction

### Xenobiotic sensor and ligand specificity

Pregnane x receptor (PXR), also known as nuclear receptor subfamily 1, group I, member 2 (NR1I2; ZDB-GENE-030903-3), is a ligand-activated zinc finger transcription factor that in mammals regulates transcriptional responses to drug and xenobiotic exposure including induction of the quintessential xenobiotic metabolizing enzyme cytochrome P450 3A (CYP3A) (Blumberg et al., 2008; Kliewer et al., 2002; Lehmann et al., 1998; Zhou et al., 2009). As with other NR1s, Pxr can roughly be divided into several functional domains consisting of an amino-terminal activation function (AF1) domain, DNA binding domain (DBD), ligand binding domain (LBD), and carboxy-terminal AF2. PXR is a promiscuous nuclear receptor, activated by diverse compounds including pharmaceuticals, pollutants, and endogenous compounds (Krasowski et al., 2005; Lille-Langøy et al., 2015; Moore, 2002). Although it is now clear that some teleost fish have lost *pxr* at some point in their evolution (Eide et al., 2018), PXR has been identified in most vertebrates including mammals, birds, and zebrafish, and its role as a xenobiotic sensor appears conserved despite substantial interspecies differences in ligand specificity (Ekins et al., 2008; Kliewer et al., 1998; Lehmann et al., 1998; Moore, 2002). Studies in humans suggest over half of all therapeutic drugs activate PXR to initiate biotransformation by phase 1 and 2 metabolizing enzymes, making PXR a critical regulator of drug metabolism (Lehmann et al., 1998). Studies in human and rodent models, and more recently in fish, show that PXR agonists are broadly distributed in the environment from anthropogenic sources such as sewage effluent and industrial (Bainy et al., 2013; Delfosse et al., 2015; Gräns et al., 2015; Kubota et al., 2015; Wassmur et al., 2010). These studies suggest that PXR is in the first line of the vertebrate chemical defensome (Dussault and Forman, 2002; Goldstone et al., 2006).

While there appears to be a conserved role for PXR as a drug and xenobiotic “gate keeper,” ligand specificity across species is inconsistent and requires direct testing of possible ligands to identify species-specific agonists. Notably, even among mammals PXR exhibits differences in ligand-mediated activation, for example, the human PXR agonist rifampicin does not activate rat PXR, while pregnane-16α-carbonitrile (PCN), a known rat PXR agonist, fails to activate human PXR (Igarashi et al., 2012; Tirona et al., 2004; Xie et al., 2000). Notably, however, the steroid pregnenolone (PN) is a broadly conserved ligand across vertebrates, including zebrafish (Kliewer et al., 1998; Kubota et al., 2015).

### *PXR* loss-of-function

Directed loss-of-function studies can establish the roles for a given gene and its product such as *PXR*. Studies using a morpholino oligonucleotide to transiently block *pxr* translation in zebrafish resulted in the loss of ligand-activated transcription of Pxr target genes, *cyp3a65* and *pxr* itself (an apparent auto-induction loop) (Kubota et al., 2015). Similar loss-of-function findings have been reported in mammals showing *Pxr* genetic knockouts do not activate *Cyp3a4* transcription in the presence of PXR ligands (Xie et al., 2000). Additionally, physiological abnormalities in *Pxr*-deficient rodents have been reported, including poor breeding success (Frye et al., 2014), reduced bone density, premature wearing of cartilage (Azuma et al., 2010, 2015; Konno et al., 2010), and disrupted glucose homeostasis (Spruiell et al., 2015). Studies examining natural human variants of PXR identified a single amino acid substitution (R98C) in the second zinc finger domain that eliminates PXR DNA binding and transactivation of *CYP3A4* in the presence of authentic PXR-ligands (Koyano et al., 2004). Similarly, zebrafish allelic variants expressed in COS cells revealed differences in *cyp3a* luciferase reporter expression in response to clotrimazole (Lille-Langøy et al., 2019).

To investigate the possibility that total loss of *pxr* in zebrafish would present transcriptional consequences consistent with those reported in the mammalian literature, and to determine further ligand-specific effects, we generated two genetic *pxr* loss-of-function lines in zebrafish using CRISPR-Cas9 RNA-guided nuclease (RGN). The two zebrafish lines each carry a sizable deletion in the *pxr* locus. *Prima facie* interpretation of these mutations would suggest loss-of-function, however, the mutant animals show activation of Pxr target transcription when exposed to pregnenolone, suggesting a compensatory mechanism is responsible for target gene transcription or that a functional Pxr protein is still being translated despite the presence of mutations.

## Results

The zebrafish *pxr* locus spans approximately 65kb on chromosome 9 and contains 9 or 10 exons with coding sequence contained in exons 1-9 (Fig 1A). Two different *pxr* lines were generated, each carrying a sizable deletion in the *pxr* locus.

**Figure 1.**
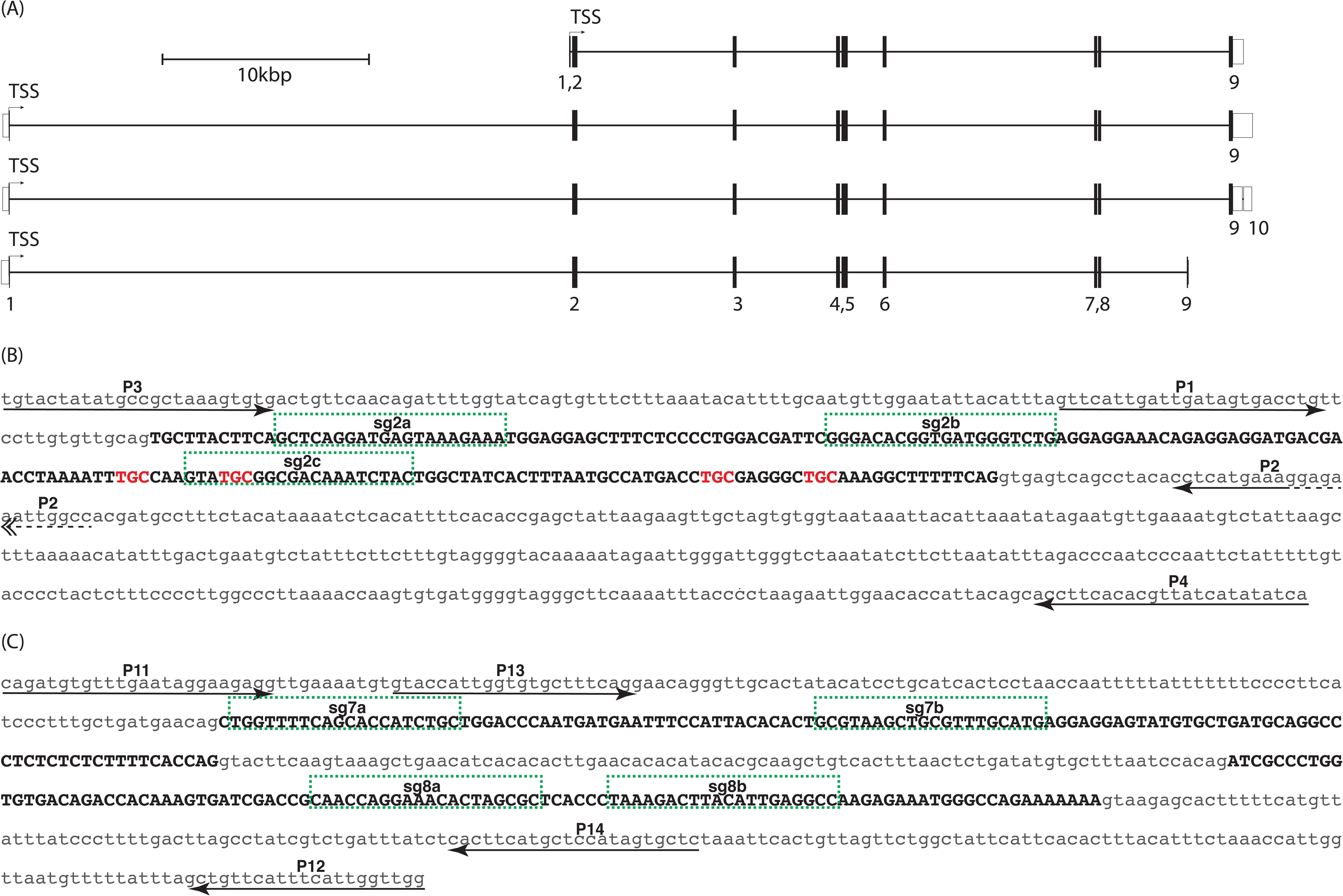
Zebrafish *pxr* gene models, sgRNA target sites and PCR primer locations. (A) Merged ENSEMBL/Havana gene models identify four transcript variants (ZDB-GENE-030903-3; Chromosome 9:9,524,490 - 9,584,607). Numbered exons are shown as open bars for untranslated sequence or solid bars for coding regions. Translational start sites (TSS) reside in exon 1 for all transcripts. Intronic regions are shown as horizontal black line spanning exons. (B) Sequence of exon 2 (black uppercase text) and adjacent intronic regions (grey lowercase text) are shown. Position and orientation of PCR primers are shown for primers P1-P4 (arrows; also see Table 2). Single guide RNA target sites are labeled, sg2a, sg2b, sg2c and outlined (green hashed boxes). Trinucleotides coding for zinc finger cysteine residues are shown in red lettering. (C) Sequence of exons 7 and 8 (black uppercase text) and adjacent intronic regions (grey lowercase text) are shown. Position and orientation of PCR primers are shown for primers P11-P14 (arrows; also see Table 2). Single guide RNA target sites are labeled, sg7a, sg7b, sg8a, sg8b and outlined (green hashed boxes).

### Exon 2 targeting

Four *pxr* splice variants have been identified in zebrafish with exon 1 contributing to a short sequence of amino acids for all transcript variants (Fig 1A) (Yates et al., 2016). Exon 2 sequence is entirely coding and conserved among the four identified transcript variants, and therefore, was selected as the primary locus for RGN targeting (Fig 1B). Pxr contains two C4 zinc fingers in its DNA binding domain, and exon 2 encodes all four cysteine residues in the first zinc finger, thought to be necessary for C4 zinc finger structures in general (Fig 1B). Thus, it seemed reasonable to predict a lesion in the zinc finger domain would compromise the ability for Pxr to bind *cis*-regulatory units and therefore abrogate its capacity for transactivation of target genes such as *cyp3a65*. Three non-overlapping single guide RNA (sgRNAs) targets in exon 2 were identified and sgRNA was synthesized (S1 Table).

### Exons 7 and 8 targeting

Exons 7 and 8 of the *pxr* locus are coding, contribute to the ligand binding domain, and are contained in all known splice variants (Fig 1A). Four non-overlapping sgRNA target sites, two in exon 7 and two in exon 8 were identified and sgRNAs were synthesized (Fig 1C, S1 Table).

### Confirmation of mutagenesis

Genomic extracts from pooled RGN-injected or uninjected sibling control embryos (5-10 embryos per pool) were isolated and probed by PCR using primers flanking the RGN target sites (Fig 1B and C). PCR products derived from uninjected control embryos migrated as a single band at the predicted wildtype length, whereas PCR product from RGN-injected embryos migrated as a molecular weight smear and a near absence of a wildtype-sized band as shown for exon 2 targets in Fig 2A. PCR amplicons from control and injected embryos were subsequently cloned into pBluescript vector (Stratagene) and Sanger sequenced to obtain sequence-specific allele information, which revealed a range of insertion and/or deletion alleles including a 108bp deletion in the *pxr^e2^* cohort (Fig 2C) and 236bp deletion in the *pxr^e7e8^* cohort (S1 Fig) compared to wildtype control amplicons. The remaining RGN-injected embryos were raised to sexual maturity as putative F0 founders.

**Figure 2.**
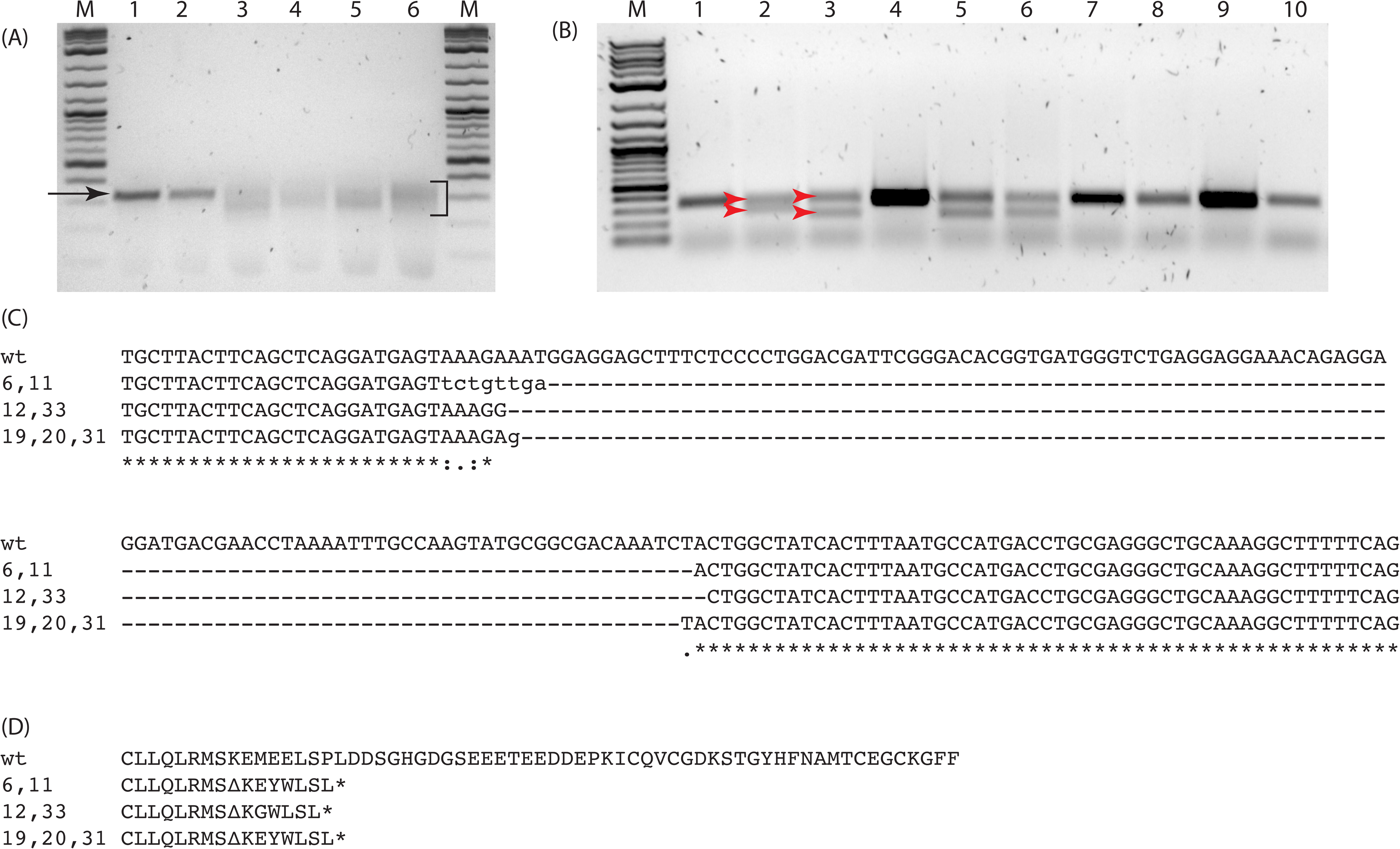
*pxr* genotyping. (A) Gel image showing PCR products derived from 24 hpf RGN-injected F0 founder or uninjected control embryos. Single band migrating at approx. 350 bp is apparent for control embryos (arrow, lanes 1,2). A smear migrating from approx. 350 - 200bp is visible for RGN-injected embryos (bracket, lanes 3-6). 2-Log DNA ladder (lanes M, NEB). (B) Gel image showing PCR products derived from F1 fin biopsies of outcrossed founder progeny (lanes 1-10). PCR products from the wildtype/deletion-mutant (*pxr^e2/+^*) heterozygotes migrate as two bands, upper and lower respectively (red arrows, lanes 2, 3, 5, 6). (C) Sequences for deletion-mutant exon 2 locus from seven F1 *pxr^e2/+^* are aligned to wt. Three mutant alleles are observed. (D) Hypothetical translation of exon2 for *pxr^e2^* alleles. Mutant alleles contain deletions (---), missense mutations (lower case text) and early translation termination codons (asterisk).

### Outcross and genotyping of F1 generation

Individual F0 putative founder adults were outcrossed with wildtype AB strain individuals to generate putative heterozygote F1 embryos. A subset of these F1 embryos was sacrificed, genomic DNA extracted, and PCR probed to confirm transmission of mutant alleles through visualization of reduced PCR product size (Fig 2B). The remaining sibling embryos were raised for approximately four weeks at which point non-lethal fin biopsies were taken from individual fish for DNA extraction and PCR amplification of the targeted region. The PCR amplicons were purified and Sanger sequenced to identify sequence-specific genotypes (Fig 2C). A variety of genotypes for each of the targeted exons were obtained including a cohort of individuals with an approximate 108bp deletion in exon 2 (Fig 2C). Computational translation of the mutant alleles showed the introduction of missense mutations and early termination codons (Fig 2D). A second cohort with a 236 bp deletion spanning from exon 7 into exon 8 (S1 Fig) was verified after (but not before) F1 outcrossing and revealed a single mutant allele. Mutant alleles from the e2 and e7e8 mutant cohorts segregated according to Mendelian principles of inheritance (data not shown).

### Response to pregnenolone in wildtype and mutant F2 larva

Individual F2 embryos from *pxr^e2^* F1 heterozygote crosses (prior to genotyping) were exposed to 3 μM pregnenolone (PN) or vehicle control (DMSO 0.05%v/v) from 48 – 72 hours post fertilization (hpf), followed by dissection, genomic DNA (gDNA) extraction and PCR genotyping (Fig 3A-E and Fig 4A). To confirm transcription of mutant alleles, RNA from a subset of F2 embryos was reverse transcribed into cDNA and PCR amplified with primers targeting exons 1 (forward) and 3 (reverse). PCR products were visualized by gel electrophoresis and showed banding patterns consistent with wildtype (wt), mutant (mut), and heterozygous (het) alleles (S2 Table; Fig 4B and C). Total RNA from wt and mut embryos was extracted, reverse transcribed into cDNA and probed by quantitative real-time PCR (qPCR) for levels of the Pxr-target genes, *cyp3a65* and *pxr* (Fig 3F and G; Fig 4C and D). Vehicle control treated wt and mut larva show similar levels of *pxr* basal transcription suggesting similar stability for mut and wt RNA, and a canonical upregulation of *cyp3a65* and *pxr* when exposed to PN (Fig 4D). A similar experiment conducted in F3 progeny of *pxr^e2^* (Fig 5), *pxr^e7e8^* (Fig 6) or wildtype cohorts (6 pools of 5 larvae per condition) also showed a canonical response to PN exposure and revealed enrichment of *cyp3a65* and *pxr* transcripts in PN but not vehicle exposed animals, consistent with previous experiments (Figs 5 and 6).

**Figure 3.**
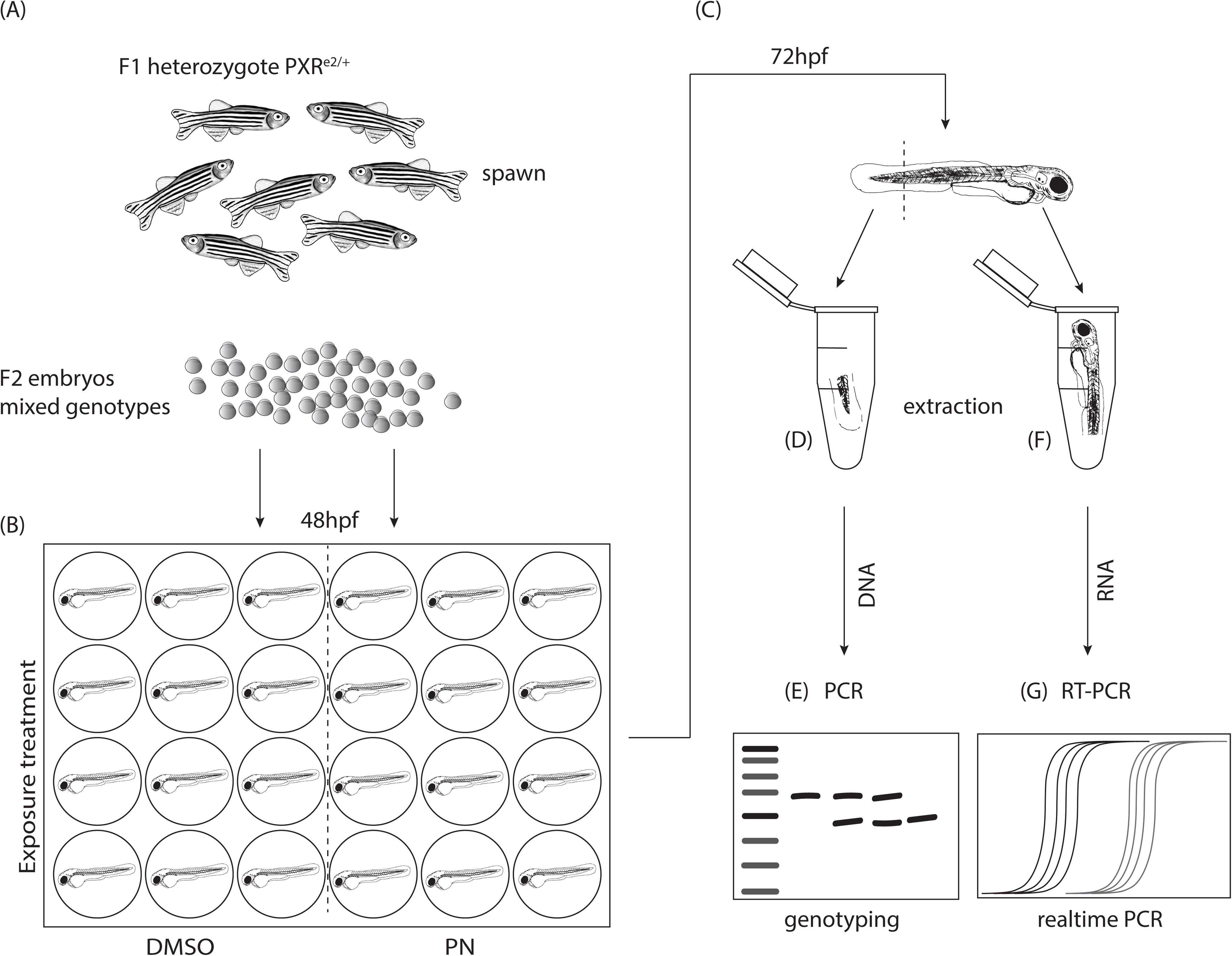
Schematic representation of experimental approach for individual embryo exposures to DMSO or PN and Pxr-target gene expression assessment. (A) Adult F1 heterozygotes *pxr^e2/+^*) were incrossed to produce mixed genotype embryos. (B) at 48 hpf embryos were transferred to 24 or 48 well plates containing 0.3x Danieau’s / 0.05%v/v DMSO or 0.3x Danieau’s / 0.05%v/v DMSO / 3 μM PN, and incubated until 72 hpf. (C) 72 hpf larvae were cut separating anterior (head/trunk) from posterior (tail) and moved to individual microcentrifuge tubes. (D) Genomic DNA isolation was carried out on tail segment. (E) PCR and gel electrophoresis was used to determine genotype. (F) Total RNA was extracted from anterior portion; (G) and reverse transcribed into cDNA for qPCR analysis.

**Figure 4.**
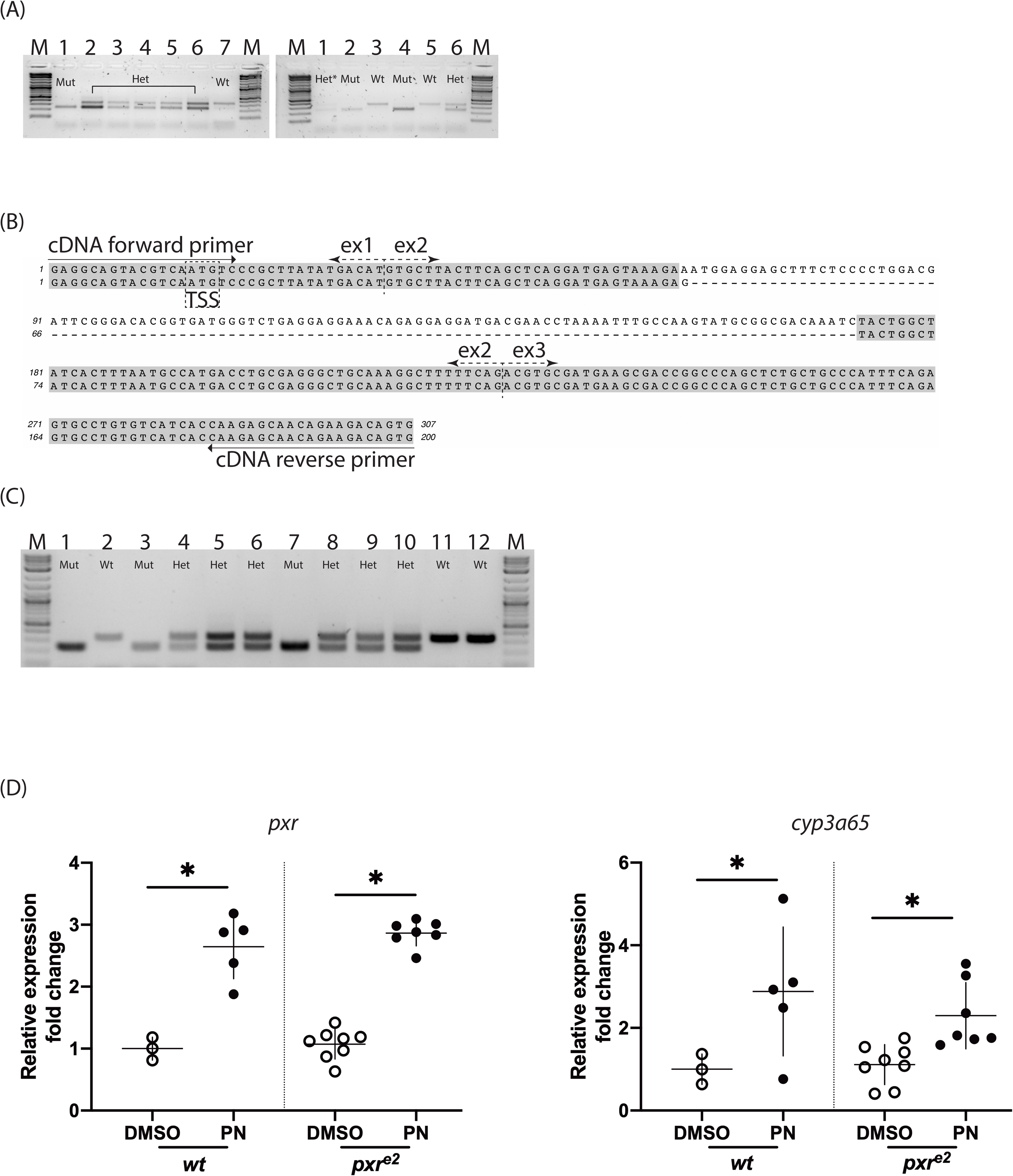
Post hoc exon 2 genotyping and gene expression of *pxr* and *cyp3a65* in 72 hpf wildtype and mutant embryos exposed to PN or vehicle control (DMSO). (A) Gel image showing PCR products derived from genomic DNA templates using P1/P2 primers (also see Figure 2, Table 2), show the presence of three product types, single lower band (Mut), upper and lower bands (Het), and single upper band (Wt). Lane marked “Het*” display weakly stained upper and lower bands. 2-Log ladder is used for size comparison (lanes M, NEB). (B) Sequence spanning exons 1-3 of transcribed mutant (bottom) allele is aligned to wildtype (top) sequence. Grey highlighted text matches wildtype sequence, whereas hashed line shows deleted region. Exon (ex), Translational start site (TSS). Primer location and orientation are shown (arrow). (C) Gel image showing RT-PCR products from cDNA primers (also see Table 2). Mut (lanes 1,2,7), Het (lanes 4-6, 8-10) and Wt (lanes 2, 11, 12) show single lower band, double band or single higher band respectively. (D) Scatter plot showing relative expression levels for the PXR-target genes *pxr* itself and *cyp3a65*. *pxr* and *cyp3a65* gene expression in individual larva are grouped by treatment (DMSO v PN) and genotype (Wt v Mut). Values are compared using 1-way ANOVA, (*P<0.05).

**Figure 5.**
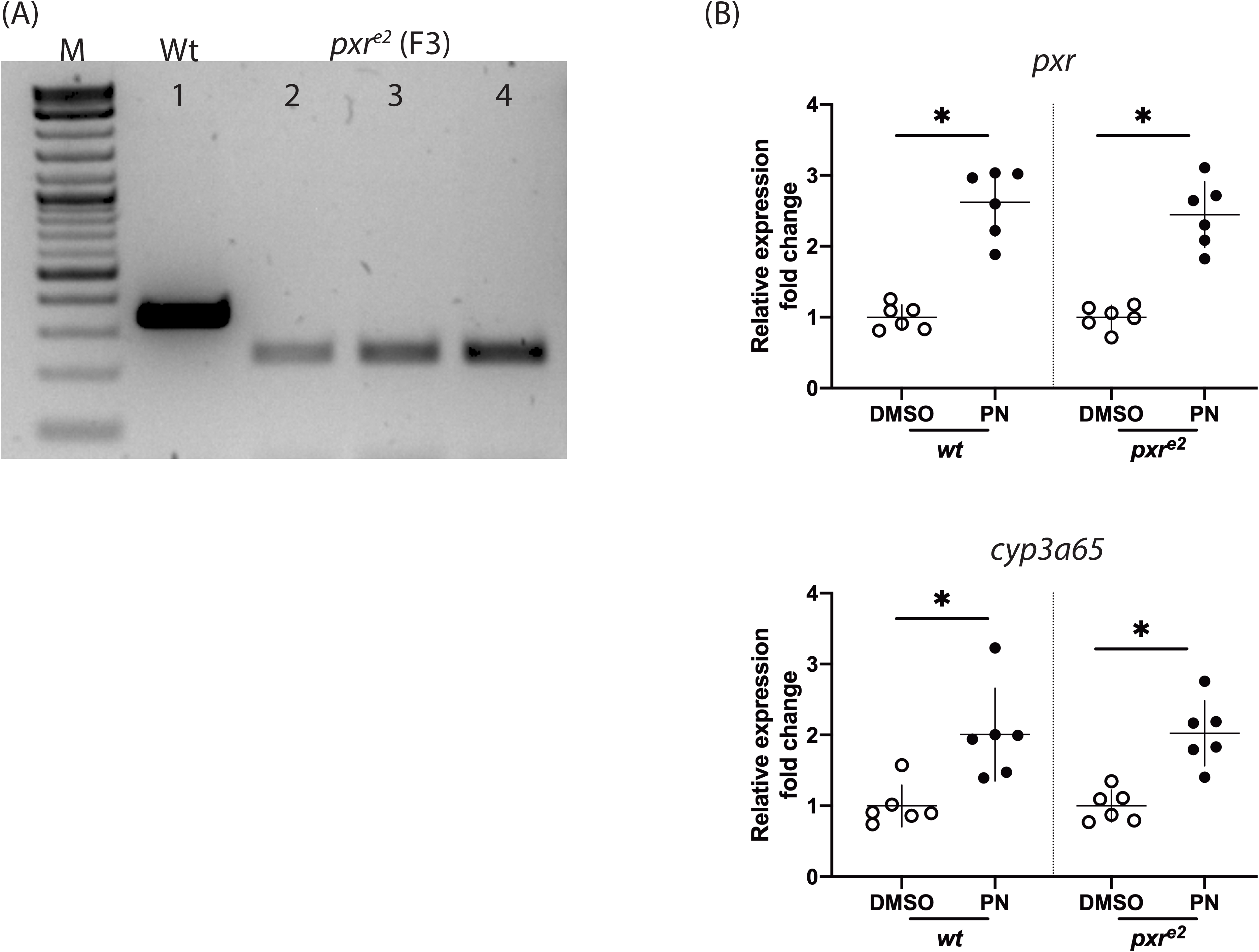
Genotype confirmation and target gene expression in *pxr^e2^* F3 larva exposed or not exposed to PN. (A) Gel image of PCR products derived from progeny of Wt (lane 1) or F2 *pxr^e2^* (lanes 2-4). (B) Relative expression of the Pxr-target genes, *pxr* and *cyp3a65*, derived from pooled larvae exposed to PN or vehicle control (DMSO) from 48-72 hpf. Each circle or dot represents one pool of 5 larvae. Expression levels are relative to vehicle treated cohort of same genotype. Values are compared by 1-way ANOVA *p<0.05.

**Figure 6.**
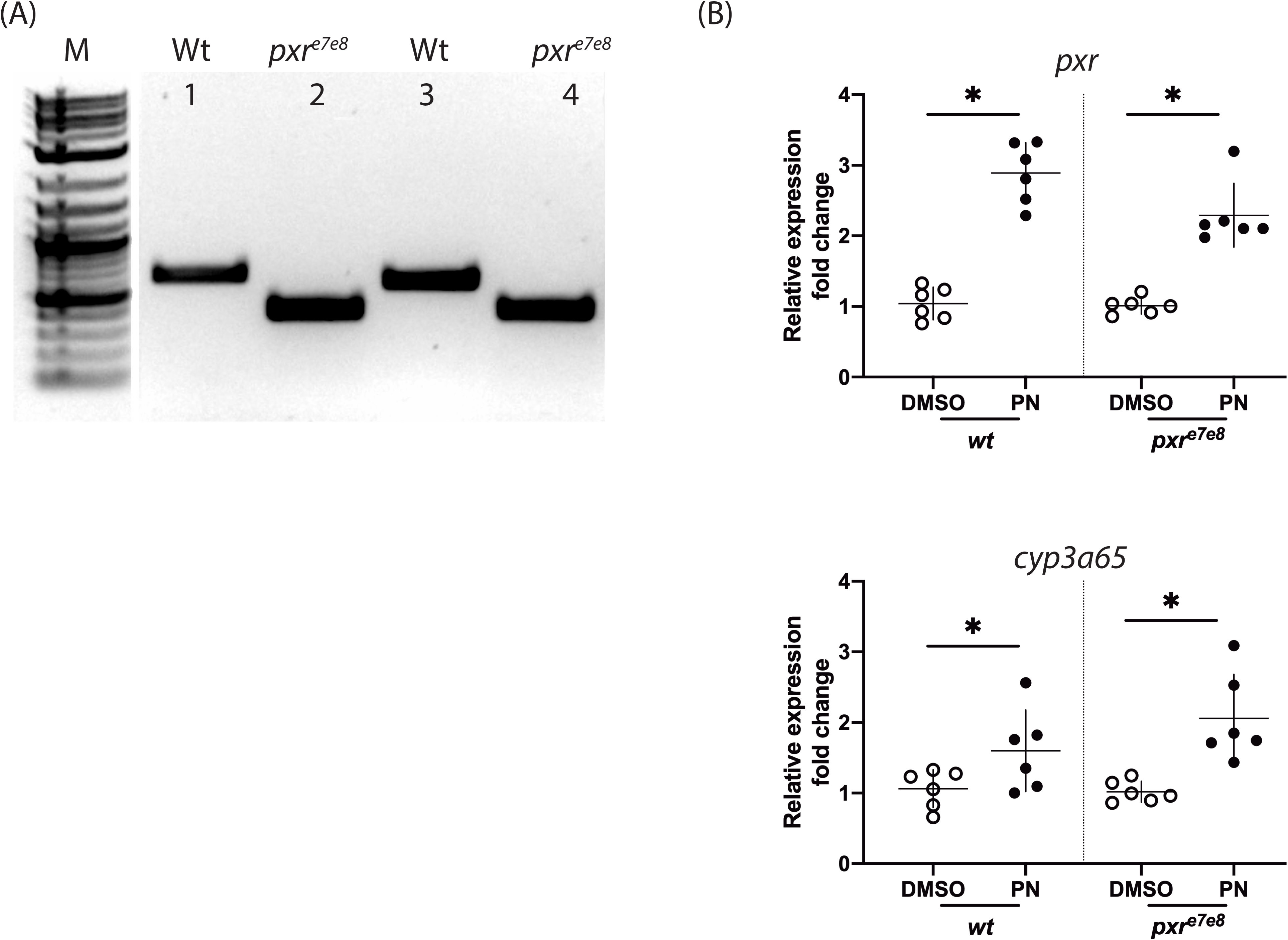
Genotype confirmation and target gene expression in *pxr^e7e8^* F3 larva exposed or not exposed to PN. (A) Gel image of PCR products derived from progeny of Wt (lanes 1,3) or *pxr^e7e8^* (lanes 2,4). (B) Relative expression of the Pxr-target genes, *pxr* and *cyp3a65*, derived from pooled larvae exposed to PN or vehicle control (DMSO) from 48-72hpf. Each circle or dot represents one pool of 6 larvae. Expression levels are relative to vehicle treated cohort of same genotype. Values are compared by 1-way ANOVA *p<0.05.

### Cloning *pxr* cDNA from mutant and wildtype fish

Total RNA was extracted from mut and wt individuals and reverse transcribed into cDNA using oligo dT priming. Primers targeting the 5’ and 3’UTRs of *pxr* mRNA were used to amplify the full open reading frame (ORF; S2 Table). Additionally, Sanger sequencing of mut full-length *pxr* ORF and subsequent translation shows the presence of a frame shift and early termination codon at the 3’ end of exon 2, however exons 1 and 3-9 appear wildtype (Fig 7).

**Figure 7.**
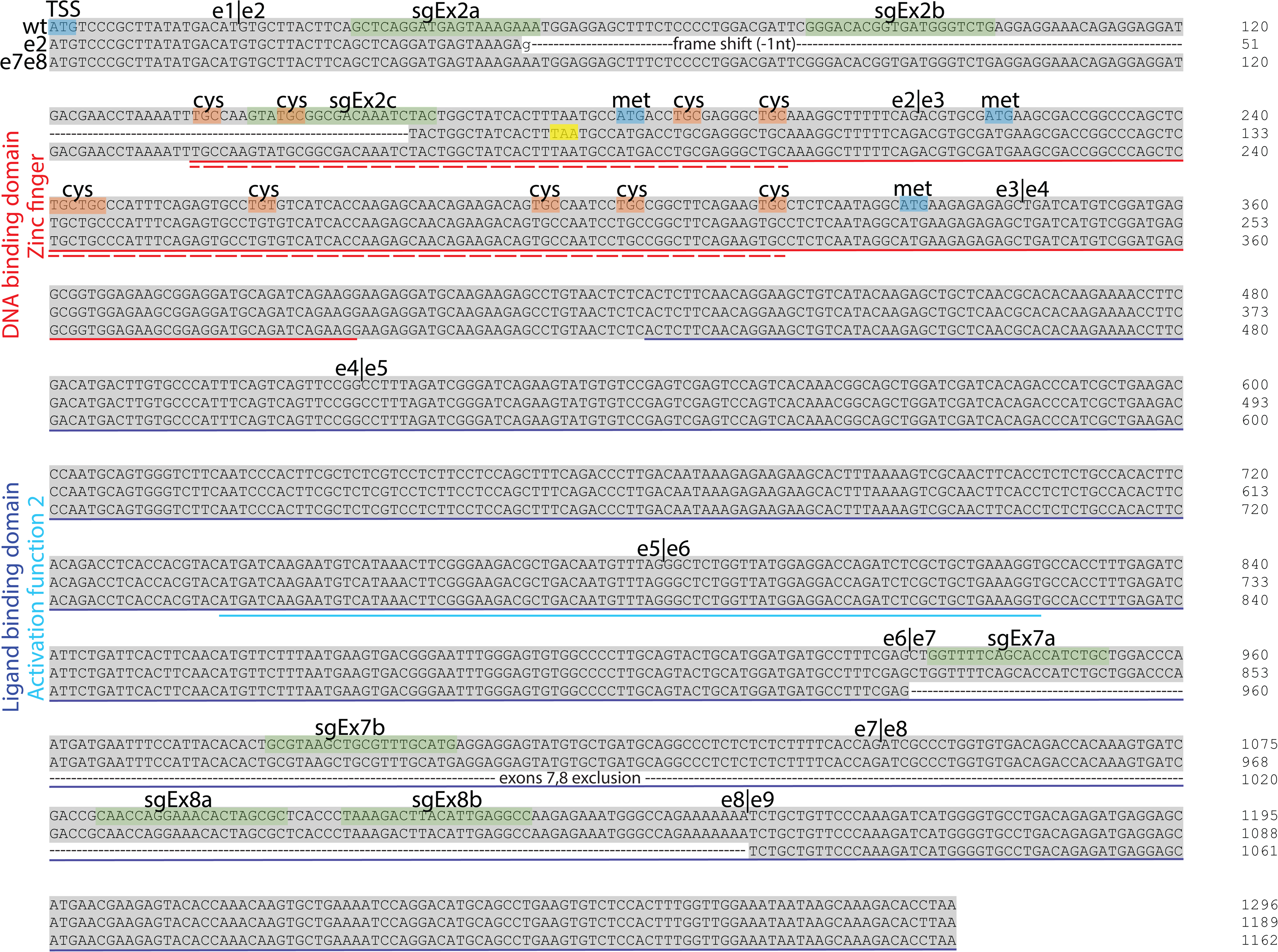
Domain map and sequence alignment of *pxr* wildtype, *pxr^e2^* and *pxr^e7e8^* cDNA. sgRNA targeting sites are highlighted in green and numbered (see also Table 1). Red underline marks putative DNA-binding domain and red dashed underline marks zinc finger domains. The C4 zinc finger cysteine codons are highlighted in orange and labelled “cys”. Blue underlined sequence marks ligand binding domain and cyan underline marks the activation function 2 domain. Exon boundaries are indicated with vertical black lines and labelled “e#”. Wildtype in frame methionine codons are highlighted in blue for exons 2 and 3 and labelled with “met”. The *pxr^e2^* mutant coding region shows 108 bp deletion and −1 bp frameshift and early *opal* termination codon highlighted in yellow. The *pxr^e7e8^* mutant cDNA shows a deletion of exons 7 and 8.

## Methods

### Animal husbandry

Experimental and husbandry procedures using zebrafish were approved by the Woods Hole Oceanographic Institution’s Animal Care and Use Committee. AB strain wildtype zebrafish were used in these studies. Embryos were obtained through pairwise or group breeding of adults using standard methods, rinsed with system water and moved to clean polystyrene petri dishes with 0.3x Danieau’s solution (Westerfield, 2007). Embryos were cultured at 28.5°C with a 14-hour light – 10-hour dark diurnal cycle. At 24 hpf 0.3x Danieau’s solution was replaced, and all dead or defective embryos were removed. Larva were fed daily with a diet according to their age starting with rotifers (*Brachionus rotundiformis)* at 5 dpf, then rotifers plus brine shrimp (*Artemia franciscana*) at 9 dpf. At 21 dpf, diets were supplemented with pellet feed (Gemma Micro 300, Skretting), and fish are fed brine shrimp and pellets only after 30 dpf. To anesthetize adult fish for fin biopsies, fish were immersed in fresh Tricaine (0.016%w/v; 3-amino benzoic acid ethyl ester; Sigma A-5040) in system water buffered with NaHCO_3_ to pH 7.5 until motionless, usually 1-2 minutes. The tip of the caudal fin was dissected using a scalpel or razor blade and processed for gDNA extraction. Following fin biopsy, adults were returned to their aquatic habitat with normal feeding regimens in place. All biopsied fish were given at least seven days to recover before any additional handling.

### Microinjection equipment

Embryos were injected using a pneumatic microinjector (Model PV-820, World Precision Instruments). Injection needles were pulled from borosilicate capillary tubes (TW100F-4, WPI) using a vertical pipette puller (Model P-30, Sutter Instruments Inc.).

### Microinjection solutions

1-2 nl of injection solution was targeted to the one-cell embryo at the interface between the embryo and underlying yolk. Injection solutions consisted of combinations of: 1 μg/μl Cas9 recombinant protein (PNA Bio, CP-01) or 400 ng/μl Cas9 mRNA (Addgene plasmid #51307 (Guo et al., 2014)) and 200 ng/μl H2B-RFP mRNA (Myers and Krieg, 2013) and pooled sgRNAs at 100-200 ng/μl for each sgRNA (S1 Table).

### Polymerase chain reaction

Endpoint PCR for genotyping or single guide RNA template preparation was carried out using Q5 (M0491 NEB) or *Taq* (M0267 NEB) polymerase and corresponding reaction buffers. Genotype PCR assembly included template (20-200 ng gDNA or cDNA), dNTPs at 200 μM final, forward and reverse primers at final concentration 300 nM (for *Taq* reaction) or 500 nM (for Q5 reaction), and polymerase-specific reaction buffer at 1x final concentration, and Q5 at 0.02 U/μl or *Taq* at 0.025 U/μl, reaction volumes range from 20 – 100 μl, and component input were scaled according to reaction size. For sgRNA template preparation proceeded essentially as described in (Bassett et al., 2013; Gagnon et al., 2014; Nakayama et al., 2014). Briefly, universal reverse primer was combined with a forward primer containing a 5’ T7 polymerase binding site, gene-specific target sequence and approximately 20 nucleotides of 3’ complimentary sequence to the universal reverse primer were combined in a 100 μl reaction at 500 nM final concentration for each, dNTPs at 200 μM final, Q5 reaction buffer at 1x final concentration, and 2 U of Q5 (S1 Table). PCR products were visualized by agarose gel electrophoresis and nucleic acid staining with SYBR safe DNA stain (S33102, Thermo Fisher Scientific), and imaged with an EZ Gel Documentation System (Bio-Rad, 1708270 and 1708273). Purification of PCR products was done using PCR QIAquick PCR cleanup kit (Qiagen, 28106), according to product instructions. Primer sequences and cycling conditions are reported in (S2 Table).

### SgRNA site selection and synthesis

Coding sequence of *pxr* exon 2 (ZFIN: ZDB-GENE-030903-3) was queried for putative targets using the web tool “CHOPCHOP” (Howe et al., 2013; Montague et al., 2014). From this, we selected three targets opting for sequences that contained a G nucleotide within the first three nucleotides of the target sequence and no predicted off-target sites. Multiple non-overlapping targets, “sgEx2 -a, -b, -c; sgEx7 -a,-b; sgEx8 -a,-b” were selected that met the aforementioned criteria (Fig 1B; S1 Table). Briefly, DNA consisting of 80-200 ng purified PCR product (see *Polymerase chain reaction - sgRNA template preparation*) was used in MEGAscript (Ambion, AM1330) or MAXIscript (Ambion, AM1309) *in vitro* transcription reactions according to product instructions with 37°C incubation lasting between 4 and 5 hours and 80 ng template DNA for MAXIscript, and 200 ng template DNA for MEGAscript reactions.

### RNA isolation

Total RNA was isolated from embryonic or larval tissue by mechanically homogenizing the tissue at room temperature in 500 μl TRIzol (Ambion, 15596-018) followed by RNA isolation according to TRIzol product instructions, or using a Direct-zol RNA MiniPrep Plus kit (ZYMO Research Corp, 2072). Contaminating genomic DNA was removed from the TRIzol isolated RNA by enzymatic digestion with 10 U of Turbo DNase (Ambion, AM2239) at 37°C for 15 minutes, in a reaction tube for TRIzol mediated extractions or on ZYMO RNA MiniPrep spin columns. DNase was removed from the RNA by organic extraction with phenol:CHCl3:IAA (isoamyl alcohol) (125:24:1) and followed with CHCl3:IAA (24:1), precipitated by adding 10%v/v 3M pH 5.2 sodium acetate solution and 2.5 volumes of 100% ice-cold ethanol and cooled to −20°C for ≥20 minutes then centrifuged at 16-20 kRCF for 20 minutes. The RNA pellet was washed two times with 70% EtOH, air dried and dissolved in 20-50 μl DNase/RNase-Free water. RNA isolated using ZYMO Direct-zol RNA MiniPrep column was eluted in 50μl DNase/RNase-Free water. Final concentration was measured by 260nm/280nm absorbance on a Nanodrop 2000. The integrity of total RNA was confirmed on a minority of samples by agarose gel electrophoresis and visual inspection of 28s and 18s ribosomal RNA bands.

### cDNA synthesis for Realtime PCR quantitation

DNA-free RNA was reverse transcribed using iScript cDNA Synthesis Kits (Bio-Rad, 170-8891) according to product instructions. RNA template amount was standardized across treatments for each experiment ranging from 400 ng for individual embryos to 1 μg for pooled samples. RNA was primed with a mix of random hexamers and oligo-dT included with the kit.

### cDNA synthesis for cloning

Up to 1 μg of DNA-free RNA was reverse transcribed using ProtoScript II Reverse Transcriptase (NEB, M0368), and anchored oligo-dT primers according to product instructions.

### Real-time PCR

Quantitative Real-time PCR was conducted on a CFX96/C1000 Realtime detector and thermocycler (Bio-Rad). iQ SYBR Green Supermix (Bio-Rad, 170-8882) or SsoFast EvaGreen Supermix (Bio-Rad, 172-5203) were used for reaction assembly as per product instructions. Real-time runs incorporated experimental templates, no template controls, and minus-RT controls, conducted with technical duplicates. ΔΔCt relative quantification was performed using Bio-Rad CFX Manager software (CFX Manager Version 3.1; Hercules, CA) normalized to zebrafish housekeeping genes, ef1a and arnt2. Cycling conditions were 95°C - 3 min; [95°C 10 sec, 60°C 30 sec]*40 cycles for iQ SYBR and 95°C - 30 sec; [95°C-5 sec, 60°C 5 sec]*40 cycles for SSoFast EvaGreen. Fluorescence was recorded during annealing-extension (i.e. 60°C). Melt curve analysis was performed over 65-95°C temperature range in 0.5°C increments and 5 sec per dwell time per step. See supplementary Table 3 for quantitative PCR primer sequences (S3 Table).

### Genomic extraction

Genomic DNA was isolated from adult fin biopsies or individual or pooled embryos 24-72 hpf. Briefly, embryos still in their chorion or single fin biopsies from anesthetized adult fish were collected and homogenized in 200 μl 10 mM Tris-HCl, 100 mM EDTA, 0.5%w/v SDS, 200 μg/ml proteinase K at 56°C for >1 hour or overnight. Homogenates were organically extracted with phenol:CHCl_3_:IAA (49.5:49.5:1) and followed with CHCl_3_:IAA (24:1). Extracts were precipitated by adding 10%v/v 3M pH 5.2 sodium acetate solution and 2.5 volumes of 100% ice-cold ethanol and cooled to −20°C for ≥20 minutes. To pellet the genomic DNA, precipitation solutions were centrifuged at 4°C and 16-20 kRCF for 20 minutes. gDNA pellets were washed twice with an equal volume of 70% EtOH, air dried at room temperature and dissolved in 10-20 μl of nuclease-free deionized H_2_O per individual embryo (e.g. gDNA from pool of 5 embryos dissolved in 50-100 μl). Final gDNA concentration was measured by 260nm/280nm absorbance on a Nanodrop 2000 spectrophotometer (Thermo Fisher Scientific).

### mRNA synthesis

1-5 μg of CS2-plasmid containing the ORF for Cas9 (Addgene #51307) or H2B-RFP was linearized by Not1 endonuclease digestion, followed by phenol:CHCl_3_:IAA extraction and EtOH precipitation. 1 μg linearized plasmid was used as template in SP6 mMessage mMachine *in vitro* transcription reaction (Ambion, AM1344) according to product instructions.

### Chemical exposure

The zebrafish Pxr agonist, pregnenolone (5-pregnen-3β-ol-20-one, PN, CAS#145-13-1), was dissolved in 100% DMSO at a concentration of 100 mM, and diluted in 0.3x Danieau’s solution for a final concentration of 3 μM PN/0.05%v/v DMSO. 48 hpf embryos were exposed as individuals or groups for 24 hours to 0.05%v/v DMSO (vehicle control) or 3 μM PN/0.05%v/v DMSO, in multiwell, plates at a volume of 1ml solution per included individual (e.g. 5 individuals in 5 mls treatment solution). Following exposure embryos (groups or individuals; whole or dissected) were snap frozen in liquid N_2_ and stored at −70°C until processing. Exposure experiments were all performed at least twice.

### Statistics

One-way ANOVA and Tukey’s multiple comparison tests were performed for gene expression data using Prism GraphPad Version 6 (GraphPad Software, San Diego, CA). Significance levels were set at P > 0.05.

## Discussion

Zebrafish are an important model organism for evaluating questions of organismal responses to toxic chemical exposure in both biomedical and ecological contexts (Stegeman et al., 2010). Integrating genome editing with toxicological studies enables mechanistic investigation into the initiating events, interactions, and adverse outcomes, advancing our understanding of the role and/or requirements of key proteins and their molecular targets in the chemical defensome (Goldstone et al., 2006). The *pxr^e2^* mutant allele lacks more than 100 nucleotides from exon 2 including nucleotides responsible for 2 of 4 cysteines in the first zinc finger domain and causes a frame shift mutation that generates premature termination codons. General cell and molecular biology principles instruct us that a genomic lesion which generates a frame shift in a coding exon should, in fact, destroy the function of the protein through missense mutations and often early translational termination.

The *pxr* gene resides on chromosome 9 with no evidence for a second copy or paralog (Yates et al., 2016). Four transcript variants are reported for zebrafish *pxr* containing 9 or 10 exons (Howe et al., 2013). In zebrafish there is no evidence for naturally occurring *pxr* splice variants lacking one or more functional domains, however in other systems, namely mammalian cell culture, alternative splicing and sequence polymorphisms result in a variety of transcript variants, including some with reduced function (Koyano et al., 2004; Lamba et al., 2005, 2004; Lin et al., 2009; Matic et al., 2010). Furthermore, Lille-Langøy et al showed zebrafish *pxr* alleles containing varying SNPs differentially induce reporter gene activation upon exposure to clotrimazole (Lille-Langøy et al., 2019). However, exposure to clotrimazole or the known human pxr ligand, rifampicin, did not induce endogenous *pxr* or *cyp3a65* expression, suggesting a possible disconnect between *in vitro* and *in vivo* studies (Salanga et al. unpublished data).

In this manuscript we report on the generation of two mutant zebrafish lines, each carrying a heritable deletion mutation in the *pxr* locus, with one affecting exon 2 (*pxr^e2^*) and the other affecting exons 7 and 8 (*pxr^e7e8^*). The *pxr* allele in *pxr^e2^* animals lacks 108 bp in exon 2 that results in a nonsense mutation. The *pxr^e7e8^* animals lack 236 bp that span from exon 7 through intron 7 and into exon 8, resulting in a frame shift and early stop codon in the transcript. We predicted that mutations would result in premature translational termination, eliminating function. In physiological terms we expected the mutations to recapitulate the loss-of-function outcomes observed from morpholino knockdown studies in zebrafish (Kubota et al., 2015) and genetic knockout studies in rodents (Xie et al., 2000), essentially abrogating canonical ligand-activated induction of target genes, such as *PXR* and *CYP3A*. Based on current zebrafish *pxr* gene models, four full-length transcripts have been identified containing identical exons 2 – 8 with variability detected in exons 1, 9 and 10. Exon 2 is the first completely coding exon for all four transcript models and contains the partial sequence for one of two zinc finger domains that comprise the DNA binding domain of the normal protein (Fig 1A). Despite the loss of two of four cysteine residues identified as components of Pxr’s N-terminal most C4 zinc finger in the *pxr^e2^* mutant, the mutant’s response to PN exposure is indistinguishable from wildtype suggesting a functional protein is being made (Figs 4 and 5). Cloning of the full mRNA from mutant and wildtype confirms the expression of the mutant mRNA, arguing against alternative splicing as a mechanism for avoiding the mutated exon 2. An important hypothesis is that translational machinery is “ignoring” the genomic lesion transcribed into the message by using an alternative start site downstream of the genomic lesion, possibly in exon 3, thereby avoiding total loss of the protein in exchange for a truncated yet functional product (Fig 8). If we assume that translational skipping is occurring and that such translation begins at the end of exon 2 or beginning of exon 3, then the resulting protein would lack one of its zinc finger (ZF) domains (Fig 8B). We would expect that this would inhibit the protein’s ability to bind DNA, which would seem a prerequisite for transactivation. Our data suggest transactivation is occurring, and therefore we surmise that a truncated Pxr is occupying *cis*-regulatory elements on target genes. We hypothesize that this may be because PXR physically interacts with RXR in the nucleus (Ihunnah et al., 2011), and thus Rxr is responsible for stabilizing Pxr binding, even in the absence of complete ZF domains.

**Figure 8.**
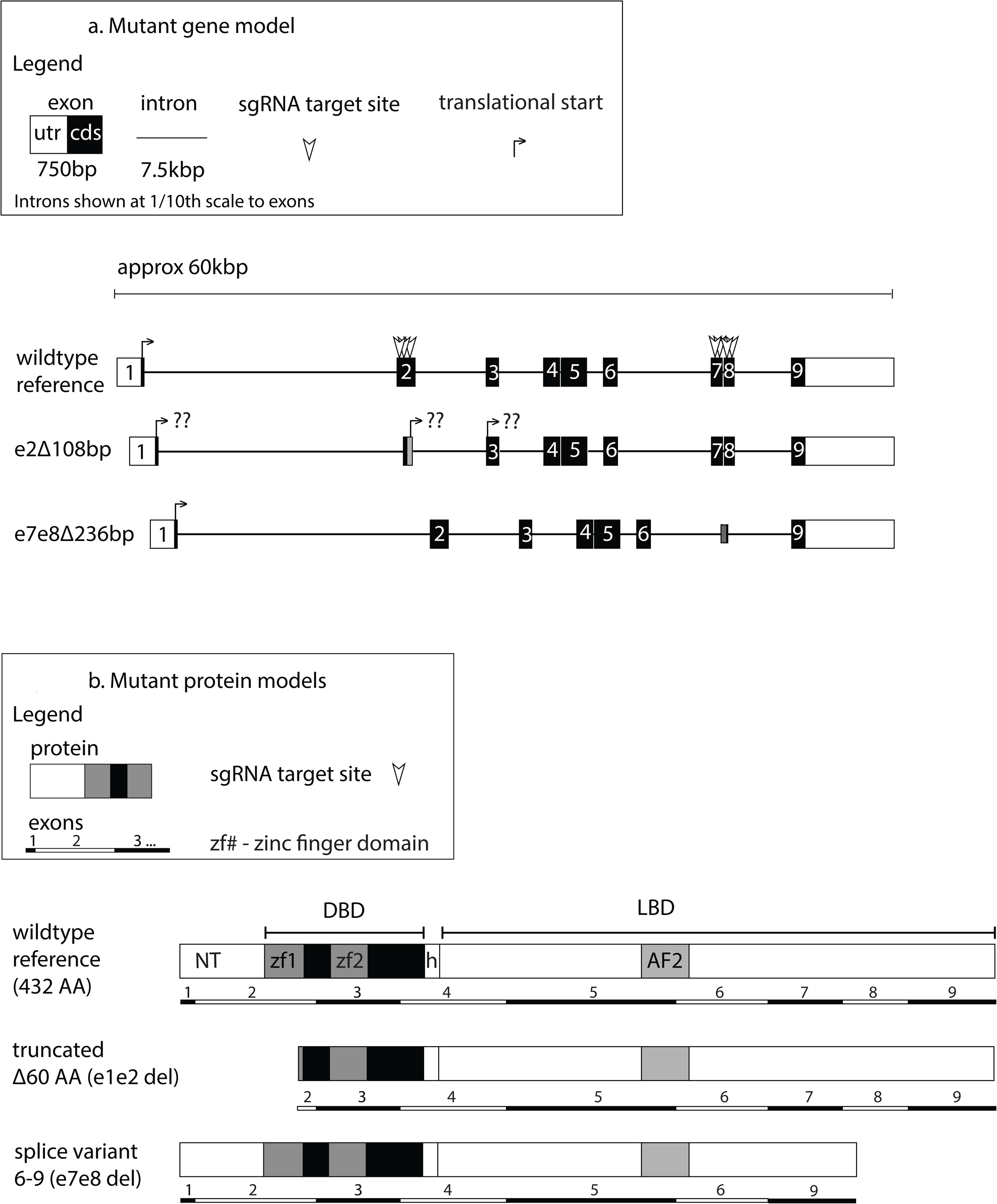
Models for gene structure and protein products for wildtype, *pxr^e2^* and *pxr^e7e8^* alleles. (A) Line diagram showing hypothetical gene model for *pxr^e2^* and *pxr^e7e8^* compared with wildtype. *pxr^e2^* model (middle) shows a shorter exon 2 and region of frame-shifted bases downstream of the lesion. Locations of potential alternative TSS according to wildtype frame and 3’ of the mutated region in exon 2 and 3 are marked (bent arrow and “??”). The *pxr^e7e8^* gene model (bottom) shows a partial deletion of exons 7 and 8 and total removal of intron 7. (B) Hypothetical protein products supposing canonical TSS of wildtype (top) and Pxr^e7e8^ (bottom) or alternative in frame translational initiation at proposed downstream TSS as proposed for Pxr^e2^ (middle). In the Pxr^e2^ model (middle) the DNA binding domain (DBD) remains mostly intact, though greater than half of leading zinc finger (zf1) has been deleted.

In the *pxr^e7e8^* mutants, cloning of messenger RNA indicated a direct splicing of exon 6 to exon 9 effectively skipping the two targeted exons (Figs 7 and 8) which if translated would result in a shortened protein. These observations point to the adaptability of transcription and translation in organisms, and highlight how genome editing technology can illuminate these processes by making normally rare events (i.e. nonsense mutations in coding exons) more common and therefore tractable for robust scientific interrogation. These results may, in fact, be an example of “translational plasticity” (Ma et al., 2019).

An alternative explanation for retaining the wildtype response to PN exposure in the mutant animals could be the induction of an alternative pathway for sensing PN and activating the downstream transcriptional response. However, there is no precedence for this scenario, and unlike mammals, teleost fish genomes do not carry a gene for the closely related nuclear xenobiotic receptor, constitutive androgen receptor (*CAR; NR1I3*). CAR has been shown to activate *CYP3A* expression and exhibit overlap in ligand sensitivity with PXR (Timsit and Negishi, 2007; Wei et al., 2002; Zhao et al., 2015). The absence of Car in fish magnifies the potential importance of Pxr for xenobiotic-sensing and response (Ekins et al., 2008), however we cannot rule out the possibility that another nuclear receptor shares ligand specificity and transcriptional activation with *pxr*, and can compensate for its loss. The morpholino effect suggests this is not the case, however (Kubota et al., 2015).

Despite all the transcriptional evidence, this study lacks direct evidence for protein translated from the mutant mRNA. No successful use of commercially available antibodies specific for zebrafish Pxr has been reported, and personal communications with researchers also investigating Pxr in fish suggest that there is a need for such a reagent. To this end, we have started efforts towards generating engineered zebrafish that have had their endogenous *pxr* locus modified to include a C-terminal epitope tag on the *pxr* coding region (N-terminal epitope tags appear to inhibit *pxr* translation; data not shown). Progress on this front will enable visualization of endogenous Pxr localization, in addition to chromatin immunoprecipitation and protein-protein interaction assays, which will provide more direct evidence for the role of Pxr in the zebrafish chemical defensome.

## Conclusion

In conclusion, our data suggest a compensatory mechanism is responsible for the PN response in zebrafish *pxr* mutants. Two alternative possibilities might also explain the outcomes: either Pxr is not required for PN-induced expression of *cyp3a65*, or a truncated yet functional mutant Pxr is responsible for the pregnenolone response. We are actively carrying out follow-up experiments to address these uncertainties including the generation of two more *pxr* CRISPR mutants, one targeting exon 3 and the other exon 6 that should effectively remove the majority of the DNA-binding or ligand-binding respectively.

## Abbreviations

AF1: Activating Function 1
AF2: Activating Function 2
CAR: Constitutive Androgen Receptor
Cas9: CRISPR associated
CRISPR: Clustered Regularly Interspaced Short Palindromic Repeats
CYP3A: Cytochrome P450 3A
DBD: DNA Binding Domain
DMSO: Dimethyl Sulfoxide
gDNA: Genomic DNA
Het: Heterozygote
LBD: Ligand Binding Domain
Mut: Mutant
NR1I2: Nuclear Receptor 1 I 2
ORF: open reading frame
PCN: pregnane-16α-carbonitrile
PN: Pregnenolone
PXR: Pregnane X Receptor
*pxr^e2^*: mutant pxr allele affecting exon 2
*pxr^e7e8^*: mutant pxr allele affecting exons 7 and 8
Rif: Rifampicin
sgRNA: Single Guide RNA
TSS: Translational Start Site
Veh: Vehicle Control
Wt: Wildtype

## Acknowledgments

We are grateful to Mark Hahn, Neel Aluru, Sibel Karchner, Diana Frank, Rebecca Helm, Anne Tarrant and Cristy Salanga for their thoughtful comments throughout this project. Support for these experiments comes from NIH P42 ES007381 and R21HD073805 (JVG).

## Supporting information

S1 Fig: genomic sequence of *pxr^e7e8^*

S1 Table: CRISPR-Cas sgRNA target and synthesis sequences

S2 Table: Primer sequences for endpoint PCR

S3 Table: Primer sequences for real-time PCR

