## Supplementary material for "CRISPR-Cas9 mutated pregnane x receptor (*pxr*) retains pregnenolone-induced expression of cytochrome p450 family 3, subfamily A, polypeptide 65 (*cyp3a65*) in zebrafish (*Danio rerio*) larvae": S1 Fig

### S1 Figure

```
wt   cctttgctgatgaacagCTGGTTTTCAGCACCATCTGCTGGACCCAATGATGAATTTCCATTACACACTGCGTAAGCTGCGTTTGCATGAGGAGGAGTATGTGCTGATGTCTTTTCACCA   120
mut  cctttgctgatgaacagCTGGTTTTCAGCACCATCTGCTGGACCCAATGATGAATTTCCATTACACACTGCGTAAGCTGC-TT------AG-----------------------------   84
     ******************************************************************************** **      **
     <--- intron 6 ---|

     GgtacttcaagtaaagctgaacatcacacacttgaacacacatacacgcaagctgtcactttaactctgatatgtgctttaatccacagATCGCCCTGGTGTGACAGACCACAAAGTGAT   240
     ------------------------------------------------------------------------------------------------------------------------   84
      |----------------------------------------- intron 7 ----------------------------------------|

     CGACCGCAACCAGGAAACACTAGCGCTCACCCTAAAGACTTACATTGAGGCCAAGAGAAATGGGCCAGAAAAAAAgtaagagcactttttcatgttatttcccttttgacttagcctatc   360
     ----------------------------------------------------------------------------------------------------ccc----gacttagcctatc   100
                                                                                                         ***    *************
                                                                                          |------------ intron 8 ----------->

     gtctgatttatctcacttcatgctc                                                                                                  385
     gtctgatttatctcacttcatgctc                                                                                                  125
     *************************
     ------- intron 8 ------->
```
