## Supplementary material for "CRISPR-Cas9 mutated pregnane x receptor (*pxr*) retains pregnenolone-induced expression of cytochrome p450 family 3, subfamily A, polypeptide 65 (*cyp3a65*) in zebrafish (*Danio rerio*) larvae": S1 Fig Legend

**S1 Figure. Partial sequence alignment of *pxr* wildtype and *pxr*$^{e7e8}$ genomic sequences.** Introns are indicated by solid black lines underneath aligned sequence. Asteriks indicate unchanged nucleotides. Dashed lines indicated deleted regions. Wildtype (wt) sequence on top and mutant (mut) sequence below. Lower case sequence indicates intronic DNA whereas capitalized sequence represent exons.
