## Supplementary material for "CRISPR-Cas9 mutated pregnane x receptor (*pxr*) retains pregnenolone-induced expression of cytochrome p450 family 3, subfamily A, polypeptide 65 (*cyp3a65*) in zebrafish (*Danio rerio*) larvae": S1 Table

| **S1 Table: CRISPR-Cas sgRNA target and synthesis sequences** | | | | | |
|---|---|---|---|---|---|
| **sg-ID** | **Exon** | **Domain included** | **Genomic target □ PAM** | **sgRNA sequence (length)** | **5’oligo sequence (additional or mismatched bases in lowercase)** |
| sgEx2a | 2 | | GCTCAGGATGAGTAAAGAAA□TGG | GCTCAGGATGAGTAAAGAAA (20) | atTAATACGACTCACTATA**gGCTCAGGATGAGTAAAGAAA**GTTTTAGAGCTAGAAATAGC |
| sgEx2b | 2 | | GGGACACGGTGATGGGTCTG□AGG | GGGACACGGTGATGGGTCTG (20) | gatTAATACGACTCACTATA**GGGACACGGTGATGGGTCTG**GTTTTAGAGCTAGAAATAGC |
| sgEx2c | 2 | DNA-binding | GTATGCGGCGACAAATCTAC□TGG | GTATGCGGCGACAAATCTAC (20) | atTAATACGACTCACTATA**gGTATGCGGCGACAAATCTAC**GTTTTAGAGCTAGAAATAGC |
| sgEx7a | 7 | ligan-binding | TGGTTTTCAGCACCATCTGC□TGG | GGTTTTCAGCACCATCTGC (19) | taatTAATACGACTCACTATA**GGTTTTCAGCACCATCTGC**GTTTTAGAGCTAGAAATAGC |
| sgEx7b | 7 | ligan-binding | GCGTAAGCTGCGTTTGCATG□AGG | GCGTAAGCTGCGTTTGCATG (20) | atTAATACGACTCACTATA**gGCGTAAGCTGCGTTTGCATG**GTTTTAGAGCTAGAAATAGC |
| sgEx8a | 8 | ligan-binding | GCGCTAGTGTTTCCTGGTTG□CGG | GCGCTAGTGTTTCCTGGTTG (20) | atTAATACGACTCACTATA**gGCGCTAGTGTTTCCTGGTTG**GTTTTAGAGCTAGAAATAGC |
| sgEx8a | 8 | ligan-binding | GGCCTCAATGTAAGTCTTTA□GGG | GGCCTCAATGTAAGTCTTTA (20) | aatTAATACGACTCACTATA**GGCCTCAATGTAAGTCTTTA**GTTTTAGAGCTAGAAATAGC |
| ***Universal sgRNA synthesis reverse-oligo sequence:** 5’AAAAGCACCGACTCGGTGCCACTTTTTCAAGTTGATAACGGACTAGCCTTATTTTAACTTGCTATTTCTAGCTCTAAAAC | | | | | |
