## Supplementary material for "CRISPR-Cas9 mutated pregnane x receptor (*pxr*) retains pregnenolone-induced expression of cytochrome p450 family 3, subfamily A, polypeptide 65 (*cyp3a65*) in zebrafish (*Danio rerio*) larvae": S2 Table

| S2 Table: Primer sequences for endpoint PCR | | | | | | |
|---|---|---|---|---|---|---|
| **Primer pair** | **Primer ID** | **Location** | **Orientation** | **Sequence (5')** | **Target** | **Polymerase: cycling conditions (wildtype amplicon length)** |
| P1/P2 | seqP1 | intron 1 | Forward | GTTCATTGATTGATAGTGACCTG | exon 2 | Taq: 95ºC 3 min, 35x[95ºC 30 sec, 60ºC 30 sec, 68ºC 30 sec], 68ºC 5 min (~267 bp) |
| | seqP2 | intron 2 | Reverse | GGCCAATTTCTCCTTTCATGAGG | | |
| P3/P4 | seqP3 | intron 1 | Forward | TGTACTATATGCCGCTAAAGTGTG | exon 2 | Taq: 95ºC 3 min, 35x[95ºC 30 sec, 60ºC 30 sec, 68ºC 1 min], 68ºC 5 min (~816 bp) |
| | seqP4 | intron 2 | Reverse | GATAACATGATAAATGCCTTCAGG | | |
| P5/P6 | seqP5 | mRNA-exon 1 | Forward | GAGGCAGTACGTCAATGTCC | exon 2 | Taq: 95ºC 3 min, 35x[95ºC 30 sec, 58ºC 30 sec, 68ºC 30 sec], 68ºC 5 min (~307 bp) |
| | seqP6 | mRNA-exon 3 | Reverse | CACTGTCTTCTGTTGCTCTTGG | | |
| P7/P8 | orfP7 | mRNA-UTR | Forward | TCAGGACTGTAACTCTGTGTTCG | ORF | Q5: 98ºC 3 min, 35x[98ºC 15 sec, 58ºC 15 sec, 72ºC 1 min], 72ºC 5min (~1457 bp) |
| | orfP8 | mRNA-UTR | Reverse | GTCCTCGGATGAAGGTCAGCA | | |
| P11/P12 | seqP11 | intron 6 | Forward | CAGATGTGTTTGAATAGGAAGAGG | exons 7 and 8 | Taq: 95ºC 3 min, 35x[95ºC 30 sec, 60ºC 30 sec, 68ºC 1 min], 68ºC 5 min (~620 bp) |
| | seqP12 | intron 8 | Reverse | CCAACCAATGAAATGAACAGC | | |
| P13/P14 | seqP13 | intron 6 | Forward | GTACCATTGGTGTGCTTTCAGG | exons 7 and 8 | Taq: 95ºC 3 min, 35x[95ºC 30 sec, 60ºC 30 sec, 68ºC 1 min], 68ºC 5 min (~493 bp) |
| | seqP14 | intron 8 | Reverse | GAGCACTATGGAGCATGAAGTG | | |
