## Supplementary material for "CRISPR-Cas9 mutated pregnane x receptor (*pxr*) retains pregnenolone-induced expression of cytochrome p450 family 3, subfamily A, polypeptide 65 (*cyp3a65*) in zebrafish (*Danio rerio*) larvae": S3 Table

| S3 Table: Primer sequences for real-time PCR | | |
|---|---|---|
| **Target** | **Orientation** | **Sequence (5')** |
| Arnt2 | Forward | CACCTTTGGATCACATCTCATTG [Goldstone et al. 2010] |
| | Reverse | TCACCCTCCTTAGACGGACC [Goldstone et al. 2010] |
| Ef1α | Forward | CTTCTCAGGCTGACTGTGC |
| | Reverse | CCGCTAGCATTACCCTCC |
| Cyp3a65 (set1) | Forward | TTCTACGCCGCCTTACAGAAG |
| | Reverse | GGTCACTCAGACCTTTCTCCG |
| Cyp3a65 (set2) | Forward | TCCATCGCAGAAGATGACGAC |
| | Reverse | GTTTTCCCCATGCTGTCAACC |
| Pxr (set1) | Forward | ATAGGCATGAAGAGAGAGCTGAT |
| | Reverse | TGTTGAGGAGTGAGAGTTACAGG |
| Pxr (set2) | Forward | ATAGGCATGAAGAGAGAGCTGAT *(same as in set 1)* |
| | Reverse | GAGTTACAGGCTCTTCCTGCATC |
